# Unveiling the neuro-vascular interplay in the skeletal muscle in health, injury and disease

**DOI:** 10.1101/2025.08.12.669843

**Authors:** Sandra Fuertes-Alvarez, Alex M Ascension, Noelia Pelaez-Poblet, Amaia Elicegui, Susana Gonzalez-Granero, Jose Antonio Gomez-Sanchez, Rosario Osta, Jose Manuel García-Verdugo, Sonia Alonso-Martin, Ander Izeta

## Abstract

Neuromuscular junctions (NMJs) are complex multicellular structures that convey motor neuron-induced responses in the skeletal muscle. Their cellular composition is well characterized, but interactions between the different cell types and with the surrounding microenvironment remain underexplored. Here, by using a panel of newly discovered mouse cell lineage markers, single-cell RNA sequencing analyses and iDisco tissue clarification, we demonstrate the existence of contacts between adjacent NMJs through their kranocytes, assembling kranocyte network-like structures within the muscle interstitium, which is essential for regulating blood flow and supporting muscle regeneration. Indeed, kranocytes do contact with the surrounding microvasculature as well, both of them uncovering a previously unexplored, expansive interactive system. Moreover, unlike the robustness shown by their counterparts, the terminal Schwann cells, we strikingly observed that kranocytes rapidly detected stressful conditions, leading to structural changes and even the loss of their connections in response to acute damage, such as muscle injury or neuromuscular disease-induced denervation. Therefore, we propose kranocytes as the interactive sensory platforms of the NMJs, working in a perpendicular axis of nerve-transmission, as key NMJ-connectors by, interconnecting adjacent NMJs and interacting with the microvasculature and the microenvironment, and also as NMJ-sensors by detecting microenvironmental inputs. Together, our data postulate kranocytes as triple connectors of the nervous, musculoskeletal and vascular systems.

## INTRODUCTION

The study of voluntary body movement transmission at motor endplates has a long history ^1^. Neuromuscular transmission occurs at the neuromuscular junction (NMJ), a complex multicellular structure through which motor neurons (MN) innervate skeletal muscles ^2^. The organization of the NMJ and the molecular events occurring at this specialized synapse are relatively well understood, and its disruption presents profound medical implications, as exemplified by a number of NMJ-related neuromuscular syndromes ^3^. Further, in some neurodegenerative diseases, such as Amyotrophic Lateral Sclerosis (ALS), muscle-driven retrograde nerve degeneration could serve as an initiation trigger well before the onset of disease ^4,5^, which constitutes the so-called “dying-back” hypothesis. Besides the MN terminal and the muscle fibre, the known NMJ structure includes two NMJ-capping cell types, which differ in origin, morphology and function: the terminal Schwann cells (tSCs, also known as perisynaptic Schwann cells) ^6^ and the kranocytes (or perisynaptic fibroblasts) ^7^. tSCs are non-myelinating Schwann cells (nmSC) that protect the synaptic area, detecting and modulating NMJ synaptic activity ^8^. tSCs are also essential for the establishment and maintenance of the NMJ structure and function during development, as well as adult NMJ remodelling and reinnervation upon injury ^8–10^. Kranocytes are CD34+ mesenchymal cells that overlie the tSCs, and extend long cytoplasmic sprouts beyond the endplates ^7,11–15^. The precise embryonic origin of the kranocyte cell lineage is unknown and their only known function is facilitating the formation of tSC bridges between adjacent motor endplates upon muscle injury, thus facilitating initiation of the endplate reinnervation process. ^7^.

Indeed, studies on the NMJ microenvironment and its potential impact in nerve transmission are limited. The interstice of skeletal muscle is tightly packed and accommodates several cell populations such as fibroblasts, fibro-adipogenic precursors (FAP), muscle satellite cells (MuSC), vascular components and immune cells in a relatively small three-dimensional (3D) space ^16,17^. Fibroblasts are a heterogeneous and versatile cell population: they produce extracellular matrix (ECM) and are able to respond to injury, promoting tissue repair and fibrosis. Specifically, recent studies link muscle fibroblasts to pro-regenerative roles following injury, identifying CD34+ fibroblasts as promoters of angiogenesis in skeletal muscle after ischemia ^18,19^. Furthermore, a muscle-resident Pdgfra+Hsd11b1+Gfra1+ mesenchymal cell population, related to FAPs, has recently been associated with the NMJs and shown to expand upon nerve transection injury ^20^. However, whether CD34+ kranocytes are part of these regenerative populations or how they interact with them is not yet understood.

The interstice of the skeletal muscle presents a highly complex network of capillary-like microvessels in-between the muscle fibres ^21,22^. The spatial relationship between endplates and capillaries has been mapped, and found to vary with muscle region and with location in the microvascular network ^23^. However, the potential crosstalk between NMJs and neighbouring microvessels remains virtually unstudied. Of note, a relevant interplay between tissue macrophages and NMJs has been described during reinnervation upon nerve injury and under pathological conditions ^24–29^. Therefore, it seems plausible that distinct cell components of the NMJ microenvironment may influence both NMJ function in homeostasis as well as NMJ reorganization upon muscle injury.

In the present study, by using a wholemount (WM) tissue clarification-based experimental approach, we unravel the existence of a new axis of communication at the NMJ level, in which kranocytes connect the NMJs with the immediately surrounding microenvironment. On the one hand, their long processes interconnect adjacent NMJs and also NMJs with capillary microvessels. On the other hand, kranocytes sense microenvironmental signals, rapidly modifying their structure, even when positioned at a distal position to the damage. Furthermore, we uncover a differential response of tSCs and kranocytes facing different types of injuries at the early stages, indicating distinct roles in sensing stress signals. Finally, we uncover cellular restructuring of tSCs in ALS and characterize for the first time the behaviour of kranocytes in this pathology. Together, our results demonstrate that NMJs interact with one another and communicate with their microenvironment providing a tissue-wide, fast-response mode to injury.

## RESULTS

### Discovery of novel markers allows fine structural characterization of NMJ-capping cells (tSCs and kranocytes) and assignment of transcriptomic cell identities

Most studies use standard tissue sections to detect mouse tSCs based on the expression of calcium-binding protein S100β, which is also expressed by myelinating Schwann cells (mSCs) (Fig. 1a). More recently described tSC-specific markers such as neuron-glia antigen 2 (NG2) ^30^ or transcription factor Tbox21 ^31^ are rarely used. In order to find potential novel specific markers of tSCs, we first reanalysed publicly available single cell RNA sequencing (scRNAseq) data of muscle interstitial cell populations ^32^. Our reanalysis of this dataset yielded two S100β+ Schwann cell (SC) clusters (Fig. 1b) in the skeletal muscle, corresponding to mSCs and tSCs (Fig. 1b’). Of note, tSCs specifically expressed the Melanoma Cell Adhesion Molecule (*Mcam*) mRNA (Fig. 1b’’ and Supplementary Table 1, highlighted in yellow), while mSCs specifically expressed genes involved in the myelination process, such as *Mpz* and *Prx* (Fig. 1b’’’) ^33–36^. MCAM (also called CD146) is an adhesion molecule highly expressed by endothelial cells ^37,38^, however, in cultured SCs, MCAM promotes cell adhesion and migration ^39^, and *in vivo* analyses showed that SCs upregulate MCAM upon peripheral nerve injury, as well as in altered myelination conditions ^40,41^. Thus, to corroborate that tSCs physiologically express MCAM at the protein level, we examined skeletal muscles from hindlimbs of WT adult C57BL6/J mice. To characterize the 3D microenvironment of the NMJs, 2-3 mm muscle pieces were tissue-clarified and WM immunofluorescence staining performed with no further tissue sectioning. Reassuringly, MCAM antigen was found co-expressed specifically by S100β+ tSCs (Fig. 1c, d, white arrows). In contrast, mSCs adjacent to the NMJ did not express MCAM (Fig. 1c, d, pink arrows), and terminal nerves were not accompanied by MCAM+ cells before reaching the endplate (Fig. 1e, yellow arrows). Transmission electron microscopy (TEM) analysis and anti-MCAM immunogold labelling confirmed the existence of MCAM+ tSCs over the terminal nerve (Fig. 1f, dotted pink line). Hence, we postulate MCAM as a novel marker for tSCs in the skeletal muscle, and which is not expressed by adjacent mSCs at the NMJ site. Interestingly, using the MCAM antibody in tissue-cleared muscle WMs allowed to visualise cytoplasmic protrusions of tSCs that extended beyond the endplate, and which were not detected by the S100β antibody (Fig. 1g, g’, yellow arrows). The existence of such cellular protrusions has been overlooked by previous literature on tSCs.

**Figure 1.**
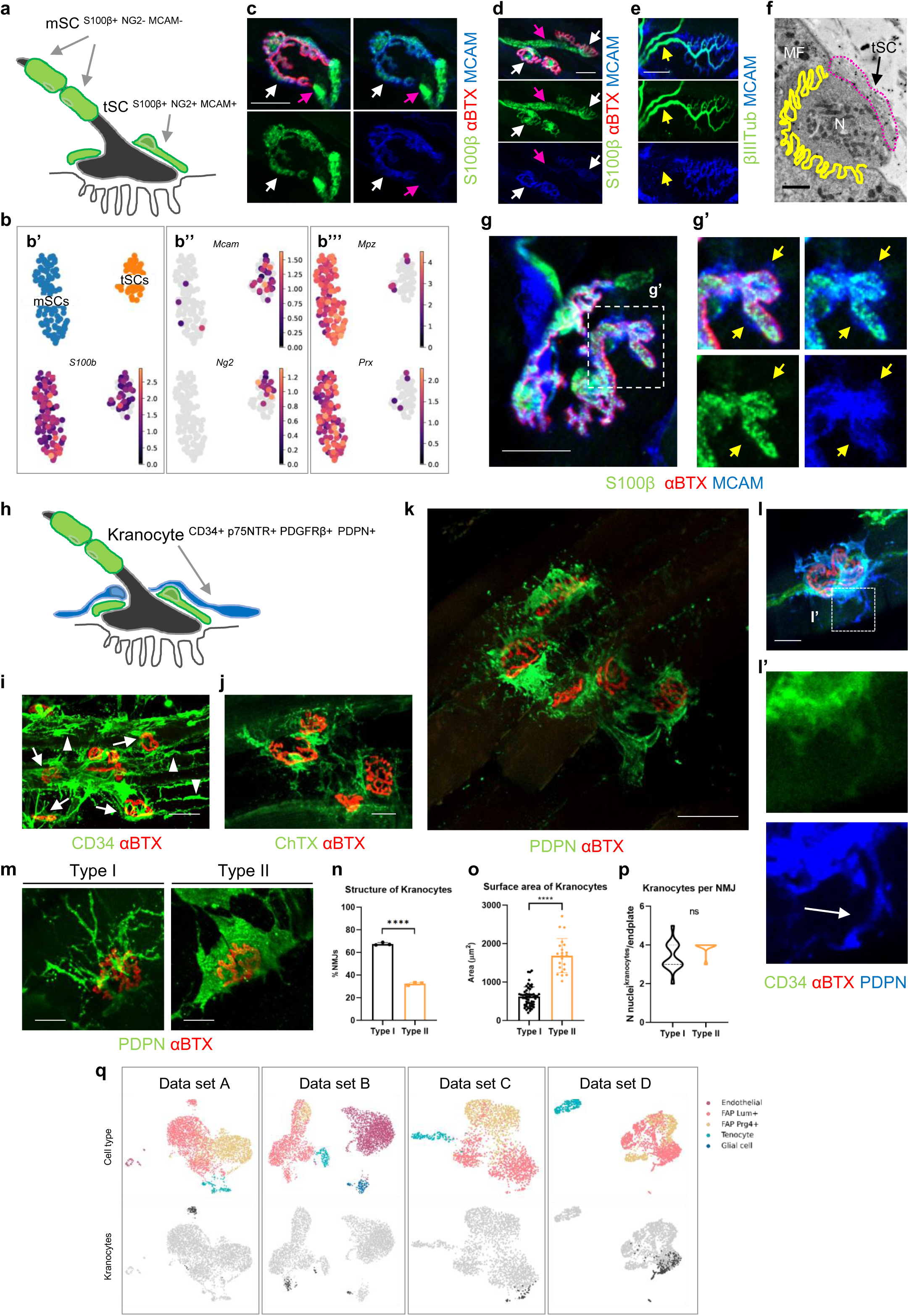
Novel markers and structural features for tSCs and kranocytes.

We followed a similar approach to further characterize kranocytes, which are CD34+ ^7^ (Fig. 1h, i, arrows). TEM immunogold analysis confirmed the presence of electron-dense CD34+ kranocytes covering the NMJs (Supplementary Fig. 1a–c, black arrows). Unfortunately, CD34 expression near the NMJ is not specific of this cell population (Fig. 1i, arrowheads): as expected, other CD34+ cells were also detected, such as interstitial fibroblasts (Supplementary Fig. 1a, yellow arrowhead) and perineural cells (Supplementary Fig. 1a, black arrowhead). Kranocytes were also highlighted by cholera toxin as previously described ^7^ (Fig. 1j), however this type of staining was difficult to combine with other antibodies. In an attempt to determine more specific markers for kranocytes, we checked a large panel of antibodies by immunofluorescence, and among them, three resulted positive in kranocytes: p75NTR (also known as NGFR), PDGFRβ, and Podoplanin (also known as PDPN): p75NTR (p75 neurotrophin receptor) is a transmembrane protein typically associated to neural lineage ^42^, PDGFRβ (Platelet-derived growth factor receptor β) is a cell surface tyrosine kinase receptor typically associated to perivascular cells ^43^, and PDPN is a transmembrane glycoprotein typically associated to lymphatic endothelium ^44^, although recently has also been associated to tumour-initiating processes ^45^, and more importantly, to nerve injury signalling ^46^. Both, p75NTR (Supplementary Fig. 1d, e, h) and PDGFRβ (Supplementary Fig. 1f, g, i) detected kranocytes, however they were quite ubiquitously expressed in muscle interstitial cells (Supplementary Fig. 1e, g, arrowheads). In contrast, PDPN marked almost exclusively kranocytes (Fig. 1k, and Supplementary Fig. 1j–l). Moreover, PDPN immunofluorescence staining revealed easier-to-detect kranocyte sprouts as compared to CD34 (Fig. 1l, l’). Thus, we selected PDPN as best available marker for kranocytes. Of note, we recognized the coexistence of two morphological types of kranocytes in homeostatic muscle, that we provisionally called Type I and Type II (Fig. 1m): the most abundant one, the Type I, was a filamentous kranocyte which partially covered the endplate (Fig. 1m, n); and the least abundant one, the Type II, was a wide kranocyte which totally covered the endplate (Fig. 1m, n). Despite the difference in structure (Fig. 1m, n) and size (Fig. 1o), there are 3-4 cells covering each NMJ, independently of Type I or Type II structure (Fig. 1p). As expected, no NMJs without kranocytes were detected.

To transcriptomically characterize the kranocytes, we reanalysed four published scRNAseq datasets of muscle interstitial cells (Fig. 1q, data set A ^32^, B ^47^, C ^48^ and D ^20^, upper panels). Based on the expression of *Cd34*, *Ngfr*, *Pdgfrb*, *Pdpn* mRNAs (Supplementary Fig. 1m) and others, a number of relatively small cell sub-clusters located within the FAPs group were highlighted as putative kranocytes (Fig. 1q, data set A to D, lower panels). Then, two different validation approaches were followed. On the one hand, it is known that kranocytes express the cell adhesion molecule tenascin C (*Tnc*) in their early migratory response to nerve injury ^7,12^. According to the data set C ^48^ (which included nerve injury analyses), some cells within the FAP cluster increased *Tnc* expression between D2 (D, day) and D10 after injury (Supplementary Fig. 1n), supporting the idea that kranocytes are transcriptomically included within the FAP population. On the other hand, we compared the putative cluster of kranocytes to the recently described “cluster 8 Hsd11b1+Gfra1+”, identified as an NMJ-associated cell population that expands upon nerve injury ^20^ (Fig. 1q, data set D). In this comparison, we observed an evident overlapping between kranocytes and “cluster 8 Hsd11b1+Gfra1+” (Supplementary Fig. 1o). Indeed, the cluster identified by our analysis shows a more restricted subgroup of cells (Supplementary Fig. 1o, first panel), suggesting that kranocytes are a particular subgroup included in “cluster 8”, which are able to react to nerve damage. Moreover, the differential expressed genes (DEG) for kranocytes of the analysed data sets were compared (Supplementary Table 2), confirming the expression of *Hsd11b1* and *Gfra1/2* in all of them (Supplementary Table 2, highlighted in green and orange). A Gene Ontology (GO) analysis of kranocytes revealed key roles as structural organizers (Supplementary Fig. 1p, q and Supplementary Table 3) at the NMJ niche, since they express *Lama2, Lamb1* and *Lamc1* (Supplementary Table 2, highlighted in blue), also identifiable by immunodetection (Supplementary Fig. 1r, white arrow). These data indicate that kranocytes are involved in the formation of the basal lamina covering the tSCs.

Finally, after analysing the interaction between tSCs and kranocytes, we detected that they could either present an interstitial gap between them (Supplementary Fig. 1a’, black asterisk) or could be in close contact (Supplementary Fig. 1b, c). Though previous studies observed kranocytes over tSCs by TEM ^7,13–15^, no specific interaction between tSCs and kranocytes has been described since tSCs are supposed to be covered by basal lamina over their entire surface ^49^. Interestingly, our data suggest that tSCs and kranocytes can interact in some specific points of their surface. Strikingly, a deeper analysis of TEM micrographs revealed kranocyte sprouts under the tSC body (Supplementary Fig. 2a, arrows, and a’). This observation was corroborated with orthogonal projections of NMJ images, where kranocyte sprouts were detected-between the S100β staining (Supplementary Fig. 2b, b’, arrows). This intricate interaction may indicate a crosstalk among NMJ-capping cells, detecting and sharing both external and internal signals to the NMJ.

Together, these data demonstrate that kranocytes are active components of the NMJs, keeping a tight relation with the tSCs and playing key roles as structural organizers.

### Kranocytes establish contacts with the surrounding NMJ microenvironment and microvascular network

Aiming at determining whether kranocytes interact with the NMJ microenvironment, we observed that kranocytes from adjacent NMJs were connected through their sprouts (Fig. 2a, b, dotted lines). Indeed, most of the NMJs appeared connected at least with one of its neighbouring NMJs (Figure 2c) through one or more sprouts (Fig. 2a, b, asterisks), forming networks of kranocytes suggesting key roles in MNJ intercommunication. Supporting this idea, we next determined whether kranocytes were able to contact other structures in the muscle insterstitium such as the vascular system. The fine structure of the microvasculature of the skeletal muscle is well known ^21–23^. To evaluate the proximity of the microvascular network to the NMJs and the NMJ-capping cells and detect possible interactions, we first analysed the localization of CD31+ capillaries close to the endplates by immunofluorescence and TEM. As expected ^23^, we observed the vascular network running parallel to the muscle fibres (Fig. 2d and Supplementary Fig. 2c). This network presented branching sites at the NMJ-enriched area (Supplementary Figure 2c, arrows), suggesting a strong demand of blood supply (oxygen and nutrients). Indeed, we detected several capillaries in the vicinity of the NMJs (Fig. 2d, arrows, and Supplementary Fig. 2d, arrows), with some of them next to the NMJ-capping cells and synaptic area (Supplementary Fig. 2d, blue arrows). We observed that tSCs were in close proximity to the vasculature (Fig. 2e–e’’, arrows, and Supplementary Video 1), even by TEM (Fig. 2f, f’, white dotted square, followed along consecutive images in Supplementary Fig. 3a, a’, white dotted squares). Indeed, analysis with MCAM staining revealed that the small tSC-protrusions MCAM+S100β-(Supplementary Fig. 2e, e’, arrows) were also CD31-(Supplementary Fig. 2f, f’), and occasionally, they were able to reach capillaries (Supplementary Fig. 2f, f’, arrows, g, g’, white arrows, g’’ and Supplementary Video 2). In all, more than the 80% of the NMJs had their tSCs in close proximity to the vasculature (Supplementary Fig. 2h). As previously mentioned, capillaries were detected by CD31+ staining and/or by MCAM+ staining (Supplementary Fig. 2i, white dotted lines) ^37^.

**Figure 2.**
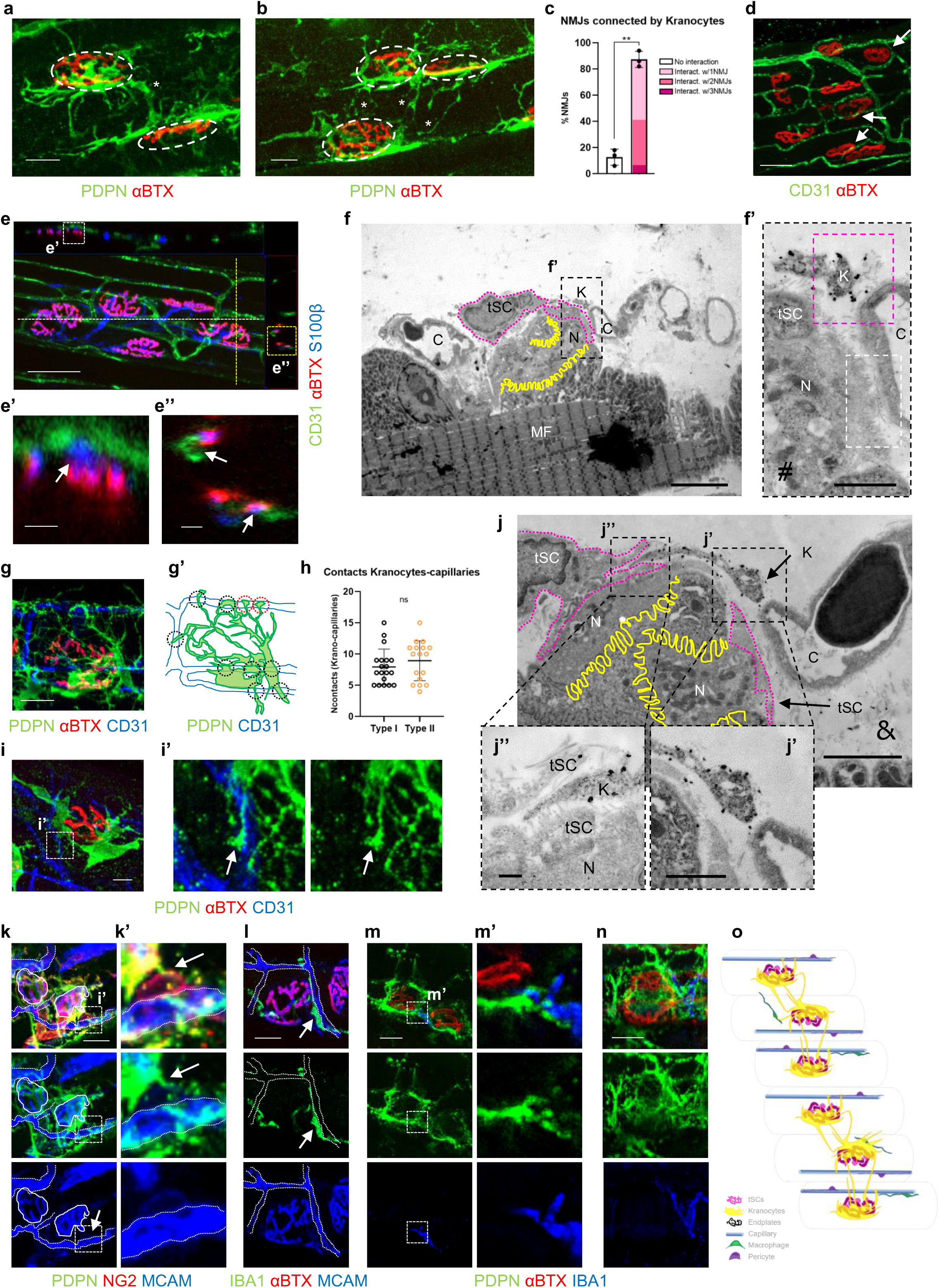
Kranocytes connect adjacent NMJs and contact NMJ-microenvironment.

Next, we interrogated whether kranocytes were able to directly interact with the surrounding microvascular network. We strikingly observed that kranocytes (Type I and Type II) exhibit multiple areas of colocalization with capillaries (Fig. 2g, g’, dotted circles, h and Supplementary Video 3). Some of these capillary-contacting sprouts reminded as end feet structures (Fig. 2 g, g’, red dotted circles, i, i’, arrows, and Supplementary Fig. 2g, g’, pink arrows and g’’). Furthermore, a detailed TEM analysis revealed sprouts from a kranocyte in close proximity to a capillary (Fig. 2f, f’, pink lined square, j, j’), both followed along consecutive images (Supplementary Fig. 3a, a’, pink dotted squares, b, b’). Part of a kranocyte under the tSC body is also observed (Fig. 2j, j’’). Additionally, our analysis revealed pericytes attached to the capillaries in the vicinity of the NMJs (Supplementary Fig. 2i–i’’), that also were contacted by kranocytes (Fig. 2k, k’, arrows). Collectively, these data demonstrate the existence of connections between NMJs and the vascular system through the kranocytes.

Finally, muscle interstitium also houses other cell types, such as macrophages. The interplay of macrophages with the NMJs during reinnervation upon nerve injury and pathological conditions has recently been described ^24–29^. Again, if there are interactions between kranocytes and macrophages has not yet been addressed. Thus, we detected the presence IBA1+ macrophages in the vicinity of the endplates (Fig. 2l–n), some of them attached longitudinally to the capillaries (Fig. 2l, arrow). These IBA1+ cells were also contacted by kranocytes (Fig. 2m, m’) or even partially covered by them (Fig. 2n). All the markers used to identify all cells at the NMJ and its microenvironment have been summarized in Supplementary Fig. 2j).

Together, these data present kranocytes as the leading connector of the NMJs with the surrounding microenvironment (Fig. 2o), bridging adjacent NMJs and contacting structures like the vascular network and other cell types such as the macrophages.

### Kranocytes respond early to denervation signals, both locally and distally, by adopting a “Primed” state

After nerve injury, tSCs form long sprouts to facilitate the reinnervation of denervated endplates ^8^. Of note, kranocytes are supposed to respond first to nerve injury and thus provide guidance to tSCs to form bridges ^7^. However, the early response (within the first day) of NMJ-capping cells to damage has never been addressed, neither their interaction with the surrounding microenvironment and vascularity. With these aims, we first evaluated the NMJ-capping cells response to nerve injury, at 1, 8 and 24h after sciatic nerve crush in both damaged ipsilateral and non-injured contralateral muscles (Fig. 3a). In tSCs, no apparent changes in cell morphology nor in the proximity of the capillaries were observed (Supplementary Fig. 4a–c’). In contrast, kranocytes, present in all endplates after nerve injury, showed an early response pattern displaying an increase of Type II kranocytes-covered NMJs already 1h after the nerve crush, and maintained in the following hours (Fig. 3b, b’). The astonishing kranocytes capacity to switch between distinct structures seemed to represent different “alert” states upon detection of stress signals. Strikingly, kranocytes early-reacted to nerve crush injury in contralateral muscles as well, by significantly increasing Type II kranocytes-covered NMJs 1h after injury (Fig. 3b’ and Supplementary Fig. 4f). These data strongly support kranocytes as key sensors of stress signals, further supporting the communication of kranocytes with the vascular system and the role of Type II kranocytes as “alert state” structures. We therefore re-named Type II kranocytes as “Primed” kranocytes, since they seem to represent a “ready to respond” stage, and Type I kranocytes as “Resting” kranocytes, since they are the most abundant type in homeostatic conditions.

**Figure 3.**
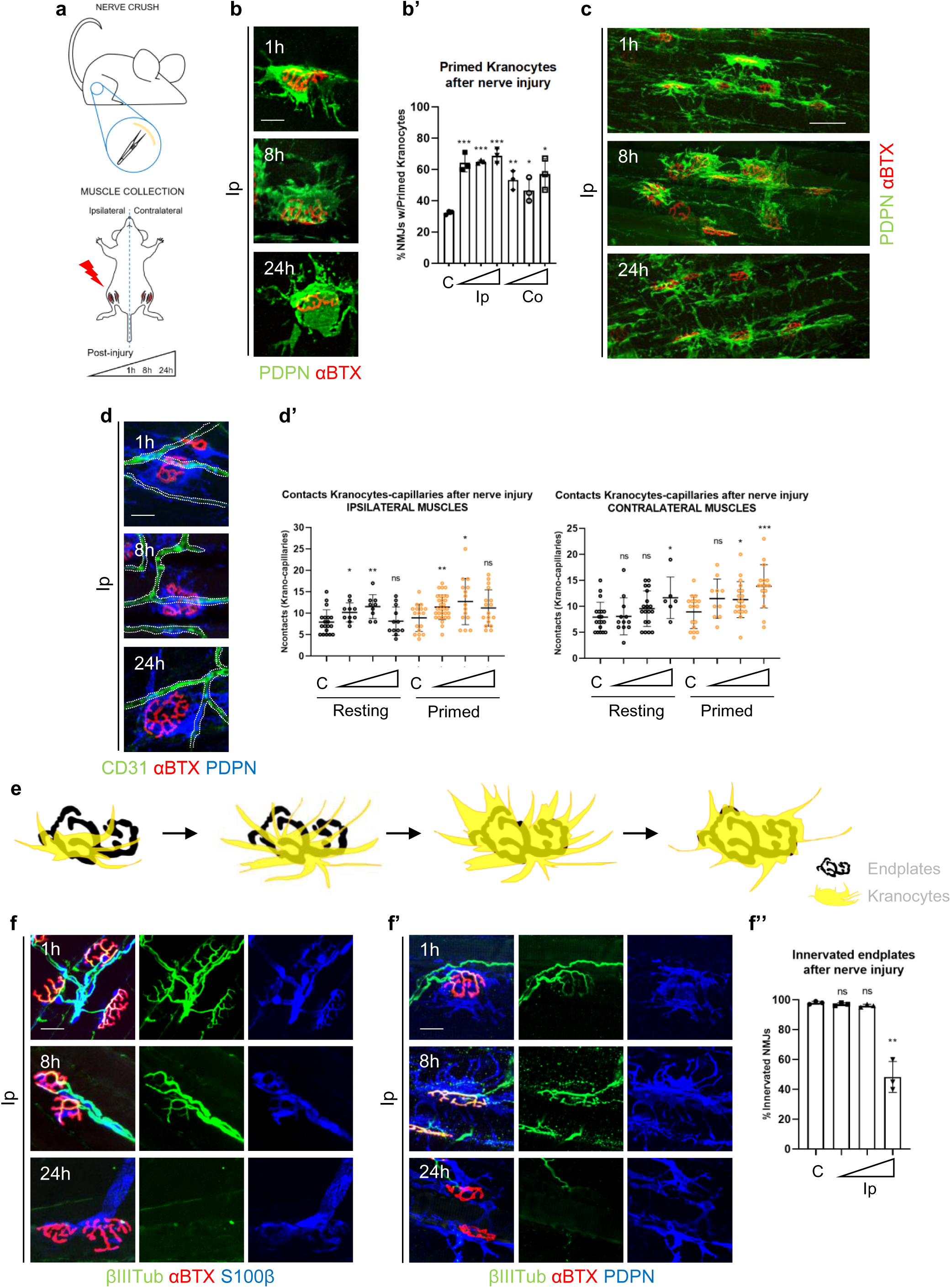
Kranocytes early-response to denervation.

Analysing the evolution of the interactions kranocytes-surrounding microenvironment, we first observed adjacent NMJs highly connected through their kranocytes (Fig. 3c), indicating that the communication network between neighbouring NMJs was still preserved after nerve damage. Moreover, quantification of the contacts between kranocytes and the surrounding microvasculature indicated that while in the ipsilateral muscles they increased 1 and 8h after damage decreasing after 24h, contralateral kranocytes increased these connections but later, starting at 8h and preserved 24h post-injury (Fig. 3d, d’ and Supplementary Fig. 4f). These data strongly support the ability of kranocytes to interact with the surrounding vascularity in order to recognize/transmit stress signals upon nerve injury, both locally and distally. In an attempt to represent both, the acquisition of the Primed state and the evolution of long sprouts (locally to the damage), a schematic representation of the possible structural evolution of kranocytes after nerve injury is shown (Fig. 3e).

Finally, we checked the retraction of nerve terminals upon nerve crush injury, observing a significant retraction 24h after injury, as terminal nerves were still in contact with the motor endplates 1 and 8h after injury (Fig. 3f–f’’). Since kranocytes were able to react from 1h after damage (Fig. 3b’, d’), this data indicates that the reaction of kranocytes is mediated by the lack of nerve transmission and not by the physical absence of the terminal nerve. These results further support the existence of a direct crosstalk between tSCs and kranocytes, where tSCs detect alterations in nerve activity and kranocytes identify the nerve malfunction through them reacting and transmitting the information to the microenvironment, the adjacent NMJs and microvasculature, activating a precautionary response, both locally and distally.

### Different responses of tSCs and kranocytes to muscle injury: the resilience of tSCs versus the acute changes of kranocytes

Since the response of the diverse cell lineages to injury in skeletal muscle is highly dependent on the injury type, we next analysed the NMJ-capping cells early response to the cobra venom cardiotoxin (CTX)-induced muscle damage by membrane depolarization ^50^. In the NMJs, CTX induces reduced cholinesterase activity and alterations in innervation ^51,52^. However, and to our knowledge, the early response of NMJ-capping cells to muscle damage has not yet been analysed. We specifically injected CTX in the *Tibialis anterior* (TA) muscle and analysed the NMJ-capping cells at 1, 8 and 24h after injury in the ipsilateral muscles [both, *in situ* (TA) and *ex situ* (rest of the ipsilateral muscles)] as well as the non-injured contralateral muscles (Fig. 4a).

**Figure 4.**
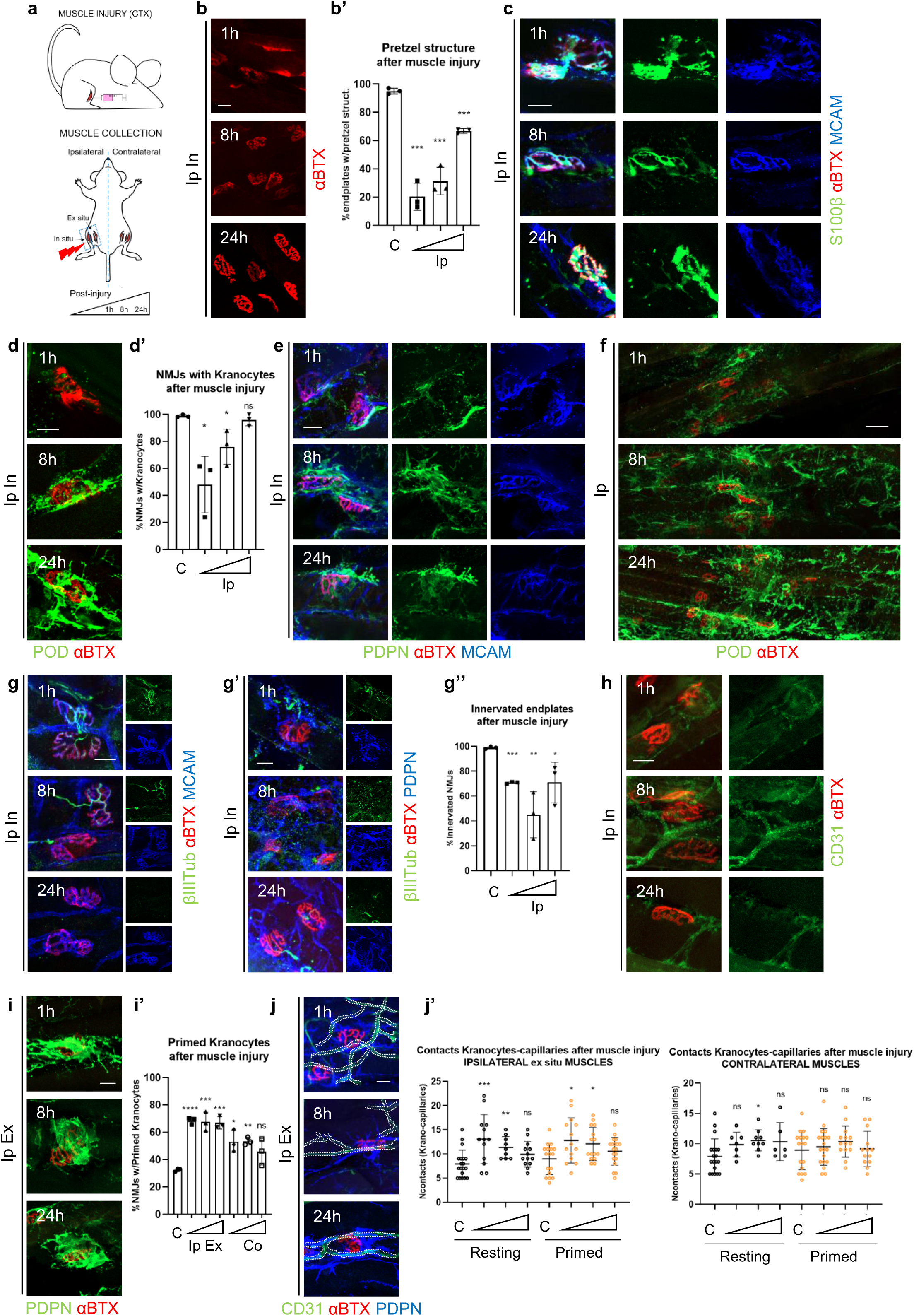
Kranocytes early-response to muscle injury.

First, the analysis of damaged muscles (ipsilateral TAs, *in situ*) showed a strong alteration of the endplates structure at early stages (Fig. 4b, b’). Despite the diffused endplates, we observed a strong resistance of the tSCs as they preserved their localization over all the analysed endplates (Fig. 4c, e). In contrast, kranocytes showed acute affection and some of them abandoned endplates during the firsts hours after muscle injury (Fig. 4d–f). Indeed, the damage was so aggressive for kranocytes that their structure was completely altered hampering their analysis (Fig. 4f). During muscle damage, even nerve retraction and abnormal innervation was observed (Fig. 4g–g’’), independently of the presence of tSCs or kranocytes (Fig. 4g, g’). These data demonstrate the different robustness of NMJ-capping cells to muscle damage, while tSCs maintain their structure and position, kranocytes rapidly react by stress conditions.

Regarding the microvasculature after muscle damage, the staining of capillaries was too diffuse for a correct analysis (Fig. 4h). Therefore, we analyse the NMJ-capping cells and their contacts with the vascularization in the ipsilateral *ex situ* muscles, adjacent to the damaged ones, and in the contralateral undamaged muscles. No alterations in tSCs structure nor in their contacts with the microvasculature were detected (Supplementary Fig. 4d, d’). However, we observed a strong reaction of kranocytes, since both, ipsilateral *ex situ* muscles and contralateral muscles, displayed a significant increase of Primed kranocytes-covered NMJs upon muscle injury (Fig. 4i, i’ and Supplementary Fig. 4g). Moreover, kranocytes connections with the vascular system increased for both Primed and Resting kranocytes 1 and 8h after damage, decreasing 24h after in the ipsilateral *ex situ* muscles, but this reaction was barely observed in the contralateral muscles (Fig. 4j, j’). Collectively, these data suggest that the acquisition of the Primed state and the formation of sprouts to contact the vascular system could follow different signalling pathways.

### tSCs and kranocytes respond differently during ischemic stress in the skeletal muscle

To determine the response of NMJ-capping cells to the lack of blood flow in the vascular system, muscle ischemia was induced by femoral artery blockage and the NMJs in both ipsilateral and contralateral muscles were analysed (Fig. 5a). While the ischemia was so aggressive that all the endplates fully degenerated 24h after injury (Fig. 5b, b’), all of them were still covered by tSCs (Fig. 5c), even when the endplate was totally unstructured (Fig. 5c, arrows). However, these tSCs presented alterations (Fig. 5c, c’), specially related to S100β staining, with dotted localization or strong decreased expression (Fig. 5c, arrows), challenging tSCs localization. Instead, MCAM revealed tSCs more efficiently, though some with altered MCAM antigen distribution (Fig. 5c, yellow arrows). tSCs managed to preserve their contacts with the vascular network during ischemia (Supplementary Fig. 4e, e’). These data further demonstrate the robustness of tSCs to survive and maintain their position under harsh stress conditions (even after endplate collapse). But also the requirement of combining the analysis of different markers, such as MCAM and S100β, to determine the tSCs presence in different contexts.

**Figure 5.**
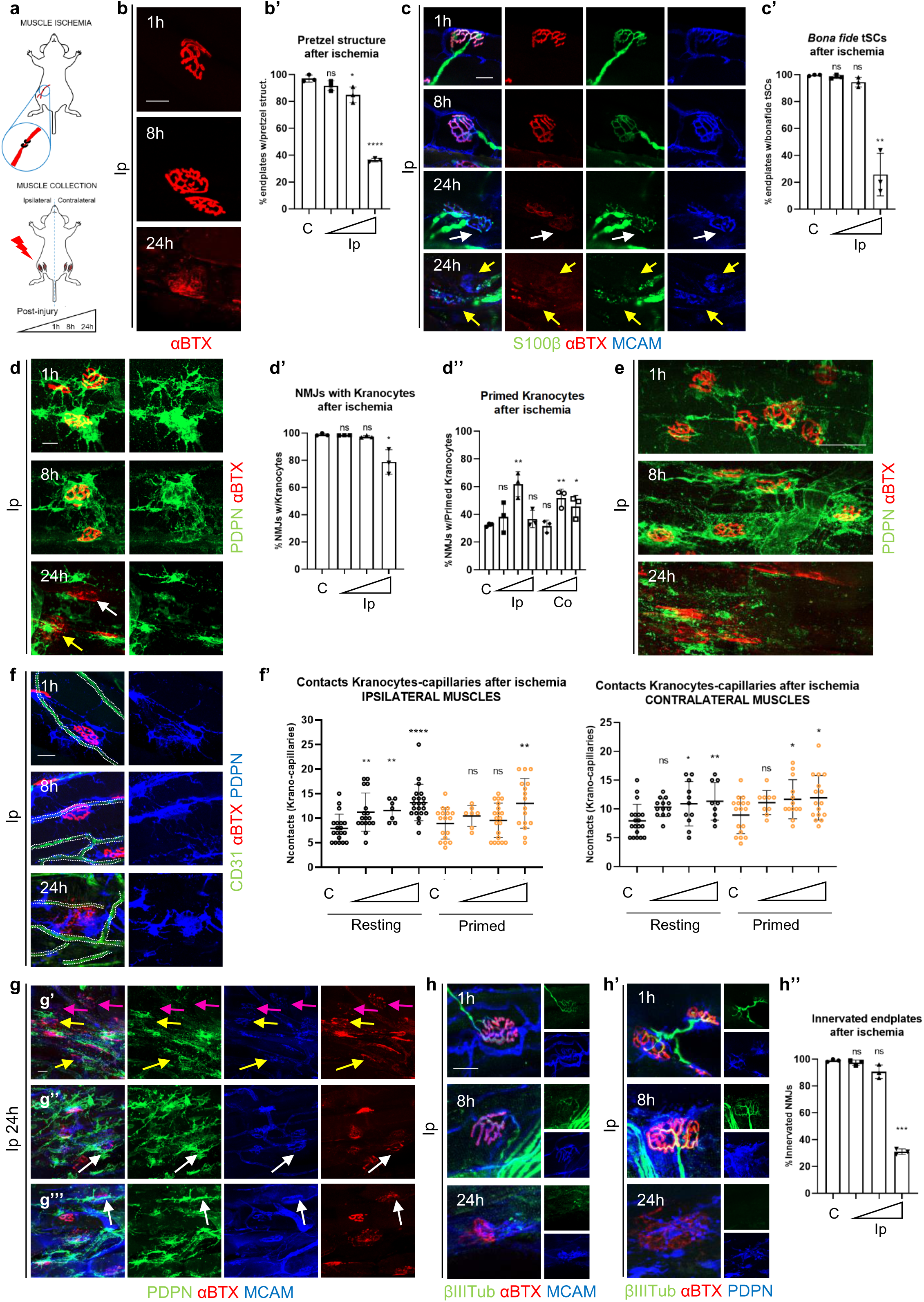
NMJ-capping cells response to ischemic signals.

Regarding kranocytes, they strongly responded to the ischemic damage with a complete altered structure 24h post ischemia (Fig. 5d, yellow arrow). In fact, more than 20% of the NMJs showed absence of kranocytes (Fig. 5d, white arrow, d’). Analysing the structure in detail, we observed that the ischemic stress increased the presence of Primed kranocytes at 8h in ipsilateral muscles but not before (Fig. 5d’’), indicating that blocking blood flow delayed kranocytes response. Even more, as ischemia persist, kranocytes collapse at 24h after damage in the ipsilateral muscles (Fig. 5d–d’’). Since blood flow was blocked, the contralateral kranocytes also delayed their response by increasing the Primed kranocytes after 8h of ischemia but still remaining after 24h (Fig. 5d’’ and Supplementary Fig. 4h). Overall, these data strongly support the kranocyte-capillary interaction sites involvement in the communication of tissue stress signals.

On the other hand, we observed that kranocytes preserve the network between adjacent NMJs until 8h after ischemia, losing the communication 24h later due to the collapse of the kranocytes (Fig. 5e). Moreover, the analysis of vascular contacts in the ipsilateral muscles revealed that Resting kranocytes seemed to rapidly react to an ischemic event (Fig. 5f, f’); however, they did not change to Primed conformation until 8h post ischemia (Fig. 5d’’). Moreover, although Primed kranocytes evidently decreased at 24h of ischemia in ipsilateral muscles (Fig. 5d’’), they increased their contacts with the vascularity (Fig. 5f, f’), further supporting the idea that the acquisition of Primed states and the formation of sprouts respond to different stress cues. Furthermore, the analysis of contralateral muscles showed that both Resting and Primed kranocytes delayed their response to the ischemic stress, since only increased their contacts with capillaries 8h after ischemia but not before (Fig. 5f’ and Supplementary Fig. 4h), reinforcing the idea of direct communication of kranocytes with the vascular system.

Finally, aiming at deeply describe the behaviour of NMJ-capping cells at 24h of ischemic conditions, we analysed tSCs together with kranocytes (Fig. 5g). On the one hand, we detected that MCAM+ tSCs were always present over the endplates (Fig. 5g, arrows), and their structure was only altered over fully collapsed endplates (Fig. 5g’, pink arrows). On the other hand, kranocytes were much more sensible to the stress, since endplates covered by tSCs displayed apparently “escapist” kranocytes (Fig. 5g’’, g’’’, white arrows), diffuse staining (Fig. 5g’, yellow arrows) or no kranocytes at all (Fig. 5g’, pink arrows). However, despite of tSCs robustness, the harsh ischemic conditions induced a strong nerve terminals retraction from their connections, even when NMJ-capping cells were still present (Fig. 5h–h’’). This result suggests that the affection of NMJ-capping cells during ischemia was so aggressive that they were unable to preserve endplate innervation.

MCAM+ tSCs altered structure over collapsed endplates (pink), scaping kranocytes (white) diffuse staining of kranos (yellow) no kranos at all (pink)

### Both tSCs and kranocytes are strongly affected in the *hSOD1^G93A^* ALS mouse model at the end stage of the disease

To further describe tSCs and kranocytes under stress conditions, we next analysed the behaviour of the NMJ-capping cells in a disease context. ALS is a devastating neurodegenerative disorder in which tSCs show inappropriate synaptic signal detection ^53^, alterations in nerve guidance after nerve injury ^54–56^ and become absent from the endplates ^57^. Interestingly, alterations in the vascular integrity of the NMJ microenvironment have been detected prior to denervation ^58,59^.

By using the *hSOD1^G93A^* ALS mouse model at end stage (>P120, or postnatal day 120), we observed an acute degeneration in the structure of *hSOD1^G93A^* endplates, with most of them displaying unstructured or diffused pattern (Fig. 6a, a’). Moreover, tSCs showed a wide variety of degeneration stages compared to WT mice (Fig. 6b, b’). As in ischemia damaged, MCAM+ tSCs were detected, but some losing the S100 staining (Fig. 6c, arrows). This striking finding revealed the existence of atypical tSCs, which modify their morphology towards a polygonal-like structure with multiple protrusions towards the muscle niche (Fig. 6b, arrows). Just 25% of the tSCs in the *hSOD1^G93A^* mice resembled the *bona fide* tSCs of the WT, while the rest presented diffuse or no S100β expression (Fig. 6b’). These atypical tSCs were not parallel to endplates, covering them partially (Fig. 6b, arrows). No endplates without MCAM+ tSCs were observed, further demonstrating that the presence/absence of tSCs cannot be exclusively determined by the detection of only S100β+ cells. Furthermore, atypical tSCs significantly increased their contacts with the microenvironment (Fig. 6c–c’’) with MCAM+ network interconnecting endplates, vascularity and other MCAM+ structures. These data demonstrate that tSCs resist well harsh conditions before leaving the endplates in ALS pathology, trying to preserve both the position over the endplates and the connections with the microenvironment.

**Figure 6.**
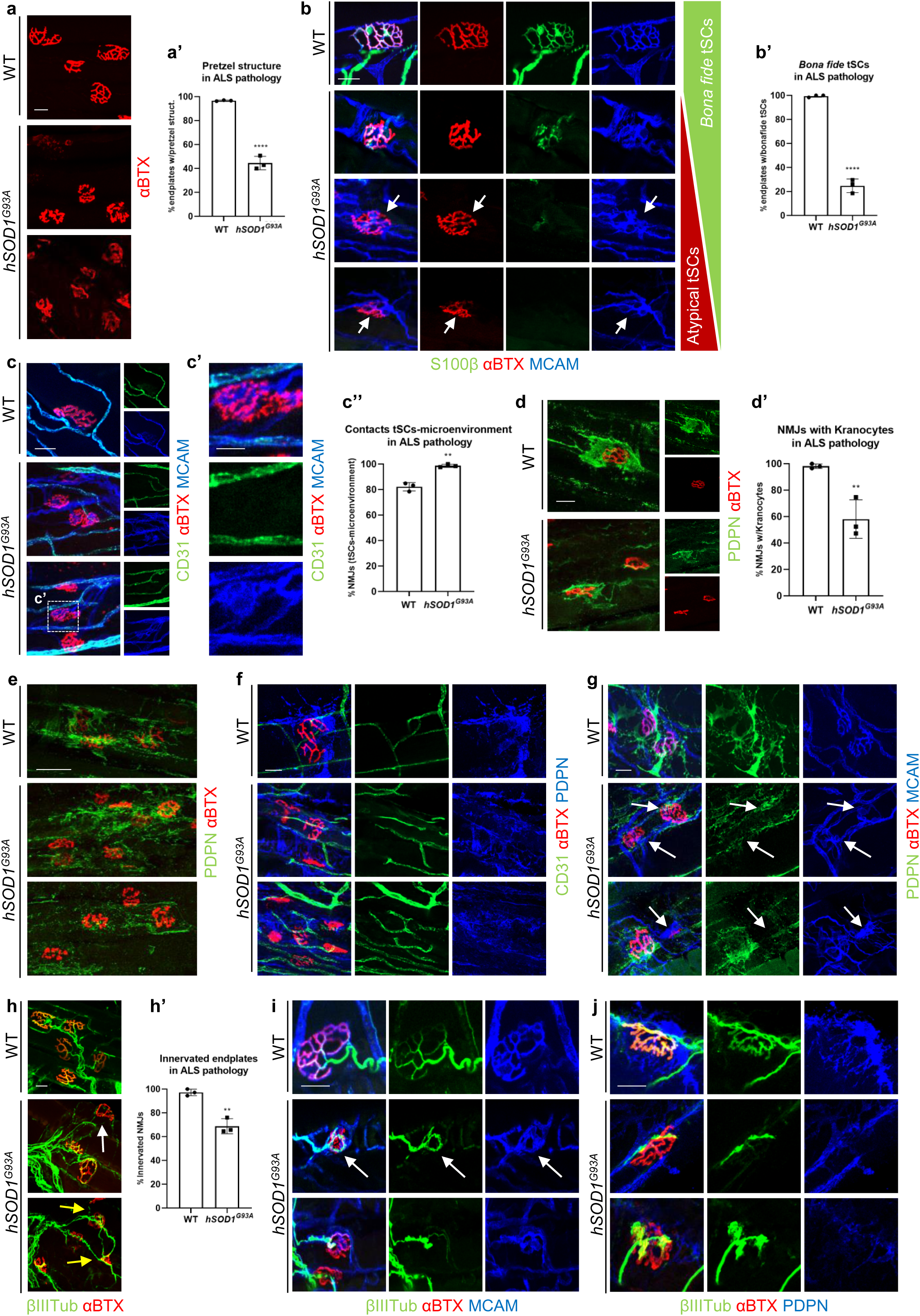
tSCs and kranocytes behaviourin ALS pathology.

Regarding kranocytes, more than 40% of the NMJs in the *hSOD1^G93A^* mice were not covered by these cells (Figure 6d, d’). Moreover, the structure of the remaining kranocytes was completely altered, making unfeasible the detection of Primed/Resting cellular states (Fig. 6d, e), nor contacts between NMJs (Fig. 6e), nor kranocyte-capillary contacts (Fig. 6f). This aberrant morphofunctional structure, that disconnect kranocytes from the NMJ microenvironment, could be disabling them to correctly respond to ALS-mediated pathology. Furthermore, when we analysed tSCs together with kranocytes, we observed that the degeneration of the NMJ-capping cells seemed to occur sequentially: the more altered tSCs, the more altered (or even absent) the kranocytes (Fig. 6g, arrows).

As denervation occurs in ALS ^60^, we also analysed the presence of terminal nerves by observing denervated endplates (Fig. 6h, white arrow) and also those with aberrant innervation (Fig. 6h, yellow arrows). A striking observation was that partial innervation was always accompanied by coverage of atypical tSCs, and endplate areas with no tSCs were denervated (Fig. 6i, arrows), strongly indicating that atypical tSCs may retain part of their role in NMJ innervation maintenance/guidance. However, innervation was observed both in the presence or absence of kranocytes (Fig. 6j), further supporting the strong alteration of kranocytes in the *hSOD1^G93A^* mice, indicating a dysfunctional phenotype in ALS pathology.

## DISCUSSION

There are several paradigms in the literature on early preventative response to distant injury or stress signals. Cardiac damage induces an early cellular response in zebrafish brain and kidney ^61^. In planarians, *Erk* waves propagate throughout body-wall muscles to induce a pan-regenerative response ^62^. In mouse MuSCs, the quiescent stem cells in uninjured (contralateral) muscle enter an “alert state” mediated by mTORC1 signalling ^63^. Some of this signalling may result in a cellular “inflammatory memory state” with broad implications in health and disease ^64^. Additionally, cellular state transitions may be regulated by changes in cellular protrusions and/or primary cilia ^65,66^, which are often mediated by signalling and/or mechanosensing pathways that induce rapid cytoskeletal reorganisation in the cell cytoplasm ^67,68^. In this article, we confirm such an injury-responsive cell state transition in the kranocytes, the outermost NMJ-capping cells of the nerve-muscle synapsis, which has not been considered before in this context. Indeed, we report the existence of “Primed” and “Resting” kranocyte states, which are regulated by their early response to different types of injury. Primed kranocytes are the most abundant type right after nerve injury, totally covering the endplate, in what could be interpreted as an attempt to protect the NMJ. However, harsh conditions such as tissue ischemia, or neurodegenerative pathologies such as ALS, led to kranocyte collapse. The data on contralateral muscles categorically showed that these cells also respond to distant injury, most likely through circulatory signalling. This is possibly mediated by the cellular long sprouts providing contacts of kranocytes with the surrounding vascular network, and supporting the distal intercommunication.

Kranocyte sprouting not only contacted the surrounding microvasculature, but also other kranocytes in the neighbouring NMJs, generating a network-like structure in the muscle interstitium supporting a previously unknown local NMJ intercommunication. In the last few years, other non-neural cells networks have been reported in different tissues such as the glial network in charge of pain sensation in the skin ^69^ or the astrocytes mesh-like network in the brain controlling neuronal activity ^70–73^. Moreover, kranocytes also showed some end feet-like contacts with the microvasculature, an interesting feature already described in brain astrocytes ^74^.

Within this work, we accurately correlate the kranocytes as a subpopulation of the Hsd11b1+Gfra1+ reactive muscle population after nerve damage ^20^ and provide data to also correlate kranocytes with the regenerative CD34+ population after muscle ischemia^19^. Moreover, future efforts will be adopted to analyse the possible correlation of kranocytes and other specific CD34+ fibroblast populations that interact with the microvasculature in different tissues, as tactocytes ^75,76^ or telocytes ^77–81^. Regarding tSCs, we have observed that they specifically express MCAM, a feature that differentiates them from mSCs. MCAM expression has been typically associated to endothelial cells and its expression is involved in the promotion of angiogenesis, permeability, and transmigration ^38,82^. Another important conclusion of this work is that only the combination of MCAM and S100β markers permitted us visualise the tSC cellular sprouts beyond the endplate. Furthermore, follow up of S100β expression to determine the presence or absence of tSCs in the NMJs is not sufficient, as we have demonstrated in ischemic conditions and in the *hSOD1^G93A^* ALS mouse model. The analysis of tSCs with MCAM revealed the remarkable robustness of tSCs during injury and disease, preserving their position over the endplates, even when the pretzel structure is degenerating, as occurs in nerve and muscle injury, in ischemia or in ALS, though losing their fitness at ALS end stages. Of note, these data challenge previous reports that claim the loss of tSCs along ALS progression, based on the analysis of the S100β marker alone ^57^.

In conclusion, here we unveil previously unknown features for both tSCs and kranocytes. On the one hand, we describe the solid robustness of tSCs facing stressing conditions through the novel MCAM described marker. On the other hand, we reveal two different roles for kranocytes, as fast-stress sensors and as participants of a complex network of local and distal communication, identifying them as the nexus point between two different and perpendicular crosstalk axes at the NMJ level: the *vertical* nerve-NMJ-muscle axis, and the *horizontal* NMJ-microenvironment-vasculature axis, acting as interactive platforms between nervous, musculoskeletal and vascular systems. Together, our data indicate the need to generate a highly descriptive mapping of these axes in order to improve different therapeutic strategies for targeted drug delivery for peripheral neuropathies and neuromuscular diseases (Stubbs, 2020).

## ACKNOWLEDGEMENTS

We gratefully acknowledge Dr. Laura Moreno-Martinez for the assistance with the maintenance of the *hSOD1^G93A^* mouse model. This work was financially supported by grants from Instituto de Salud Carlos III (ISCIII) from the Spanish Government and co-funded by the European Union (projects: PI19/00175, PI19/01621, PI22/00433, PI22/01247), and by ISCIII Programa Fortalece del Ministerio de Ciencia e Innovación (FORT23/00026); by the Elkartek Program of the Department of Economy and Competitiveness of the Basque Government (KK 2019/00093), by Diputación Foral de Gipuzkoa (2020-CIEN-000057-01, 2021-CIEN-000020-01) and by Osasun Saila, Eusko Jaurlaritzako (2020111032, 2023111035, 2023333042, 2024333017). S.F.A. was supported by a Sara Borrell fellowship from the Spanish Health Institute Carlos III (CD21/00066) from the Spanish Government. A.M.A. and A.E. were supported by the Department of Education of the Basque Country (PhD fellowships PRE_2021_2_0111, PRE_2020_1_0119). J.A.G.S. was supported by a Miguel Servet fellowship from the Spanish Health Institute Carlos III (CP22/00078). S.A.M. was supported by Gipuzkoa Fellow of Talent Attraction and Retention (2019-FELL-000010-01, 2020-FELL-000016-02-01, 2021-FELL-000013-02-01) and CIBER -Consorcio Centro de Investigación Biomédica en Red-(CB06/05/1126, Group 609; J.M.G.V. by CB06/05/1131 Group 113 and R.O. by CB21/13/00087).

## AUTHOR CONTRIBUTIONS

Conceptualization, A.I. and S.F.A.; methodology, S.F.A., A.M.A., N.P.P., A.E., S.G.G., and J.A.G.S.; investigation, S.F.A., A.M.A., N.P.P., A.E., S.G.G., J.A.G.S., R.O., J.M.G.V., S.A.M. and A.I.; data curation, S.F.A., A.M.A. and N.P.P.; formal analysis, S.F.A., A.M.A. and N.P.P.; software, A.M.A.; writing-original draft, S.F.A. and A.I.; writing-review and editing, S.F.A., J.M.G.V., S.A.M. and A.I.; project administration, S.F.A.; funding acquisition, A.I., S.A.M. and S.F.A.; supervision, S.A.M. and A.I.

## COMPETING INTERESTS

SAM is a named inventor on a patent related to neurological disorders. SAM also have ownership in Miaker Developments S.L., a startup related with a pipeline on Neurodegenerative and Neuromuscular Diseases. Rest of the authors declare no competing interests.

## MATERIALS AND METHODS

### Mouse models and procedures

All animal procedures were performed under protocols approved by the institutional committee of Biodonostia Health Research Institute, in agreement with Spanish/European regulations on animal use: PRO-AE-SS-193 (OH-20-37), PRO-AE-SS-235 (OH-21-42) and PRO-AE-SS-261 (OH-22-010). Mice were housed in the animal facility in ventilated cages under a 12/12 h light/dark cycle and with access to food/water *ad libitum*. C57BL/6J mice were acquired (Charles River Laboratories). Adult young mice were used (3-5 months) and both male mice and female mice were included in experimental groups (50/50 when possible). *hSOD1^G93A^* mouse model was purchased from JAX (*B6SJL-Tg(SOD1*G93A)1Gur/J*, Strain #:002726) and used at end stage (P120, postnatal day 120): at this stage, the *hSOD^G93A^* mice suffered from semi-paralysis. During surgical procedures, anaesthesia was induced and maintained with inhaled isoflurane, and heating plates were used to prevent animal body temperature from falling. Surgical procedures were applied just in one hindlimb (ipsilateral). After surgeries, animals recovered in their cages under supervision. Afterwards, animals were anesthetized and sacrificed by cervical dislocation (1h, 8h and 24h) and muscles from both hindlimbs (ipsilateral and contralateral) were obtained: *Tibialis anterior* (TA), *Extensor dignitary longus* (EDL), *Gastrocnemious* (Gc), *Soleus* (S).

### Surgical nerve injury

Sciatic nerve crush was performed as follows. Anaesthesia was induced in a cage with 4% of isoflurane (O_2_ flow 1l/min) until no response to stimulus (pedal reflex) was observed. Then, mouse was located on a warmed surgical table (37°C). and anaesthesia was maintained with 2% of isoflurane (O_2_ flow 1L/min) via nosecone. Animal was immobilized using surgical tape. Nerve injury was performed in one hindlimb (ipsilateral hindlimb), where hair was previously removed. Incision was performed in the outer part of the thigh from the center of the medial thigh with fine scissors (length of 1 cm approx.). Forceps and blunt scissors were used to open the incision, in order to expose the sciatic nerve. The sciatic nerve was carefully exposed and crushed three times for 15 s with three different rotation angles using angled forceps. The wound was closed using veterinary autoclips (AutoClip System). After surgery, mice recovered in their cages under supervision. Afterwards, animals were anesthetized and sacrificed by cervical dislocation at 1h, 8h and 24h post injury and ipsilateral and contralateral muscles were obtained.

### Cardiotoxin injury

Cardiotoxin (CTX) (Latoxan L8102) treatment was performed as follows. Anaesthesia was induced in a cage with 4% of isoflurane (O_2_ flow 1L/min) until no response to stimulus (pedal reflex) was observed. Then, mouse was located on a warmed surgical table (37°C). and anaesthesia was maintained with 2% of isoflurane (O_2_ flow 1L/min) via nosecone. Animal was immobilized using surgical tape. CTX was diluted in sterile 1X PBS (10µM) prior to injection. CTX injection was performed in one hindlimb (ipsilateral hindlimb), where hair was previously removed. A volume of 50 µL of diluted CTX was injected in the TA (ipsilateral TA: “*in situ*” damage; rest of ipsilateral muscles: “*ex situ*” damage. After injection, mice recovered in their cages under supervision. Afterwards, animals were anesthetized and sacrificed by cervical dislocation at 1h, 8h and 24h post injury and ipsilateral (*in situ* and *ex situ*) and contralateral muscles were obtained.

### Muscle ischemia

Muscle ischemia was performed as follows. Anaesthesia was induced in a cage with 4% of isoflurane (O_2_ flow 1L/min) until no response to stimulus (pedal reflex) was observed. Then, mouse was located on a warmed surgical table (37°C). and anaesthesia was maintained with 2% of isoflurane (O_2_ flow 1L/min) via nosecone. Animal was immobilized using surgical tape. Ischemia was performed in one hindlimb (ipsilateral hindlimb), where hair was previously removed. Incision was performed in the inner part of the hindlimb, from the center of the medial part of the femur towards the abdomen with fine scissors (length of 1 cm approx.). Forceps and blunt scissors were used to open the incision and to dissect the inguinal fatty tissue, in order to expose the neurovascular bundle. The femoral artery was carefully isolated from the femoral vein in the region distal to the inguinal ligament and proximal to the lateral circumflex branch, inserting the tip of a closed forceps between the vein and artery and create a gap between them by releasing pressure on the forceps. For limb ischemia, ligation of femoral artery was performed with 6-0 suture (polypropylene, non-absorbable), multiple knots were used. Tightness was visually checked. Skin incision was closed using the same suture. After surgery, mice recovered in their cages under supervision. Afterwards, animals were anesthetized and sacrificed by cervical dislocation at 1h, 8h and 24h post injury and ipsilateral and contralateral muscles were obtained.

### Wholemount clarification and immunofluorescence

Muscles were isolated and fixed in Histofix (Panreac 256462) O/N at 4°C in gentle agitation and stored in 1X PBS at 4°C until their use. Immunofluorescence (IF) was applied to muscles using clarification strategy as follows. Muscles were longitudinal divided to obtain small pieces of 2-3 mm of diameter approx. iDisco clarification ^83^ was applied to obtain deeper images with higher quality. A more complete 3D visualization of the NMJ region was obtain compared with common IF protocols or with muscle cryosections. The protocol used was as follows ^83^: muscles were preincubated in permeabilization solution (1X PBS with 20% DMSO, 0.2% TX100, 0.3M glycine) at 37°C O/N, washed during 24h at room temperature (RT) in wash buffer PTWH (1X PBS with 0.2% TW20, 10 µg/mL heparin), blocked with 2X PTDB solution (1X PBS with 10% DMSO, 0.2% TX100, 6% BSA) at 37°C O/N, incubated with primary antibodies (diluted in blocking solution and wash buffer 50%/50% [1X PTDB]) during 4 days at 37°C, washed with PTWH O/N at RT, incubated with secondary antibodies (diluted in1X PTDB) during 4days at 37°C (when necessary, α-bungarotoxin (αBTX) to detect acetylcholine receptors [AChRs] was included in this step), washed with PTWH O/N at RT. Muscles were also incubated with DAPI 3h at RT. Gentle shaking was used in all steps when possible, avoiding the light exposure once the secondary antibodies were used. Primary antibodies used were as follows: rat IgG2a-anti-MCAM (1:100, BD 562230), rabbit IgG-anti-S100β (1:3, Dako GA504), rabbit IgG-anti-NG2 (1:100, BD 554275), hamster IgG-anti-Podoplanin (PDPN) (1:100, Biolegend 127402), rat IgG2a-anti-CD34 (1:100, BD 553731), goat IgG-anti-PDGFRβ (1:100, R&D Systems AF1042), rabbit IgG-anti-p75NTR (1:100, Promega G3231), rabbit IgG-anti-p75NTR (1:100, Millipore AB1554), rb IgG-anti-Laminin (1:100, Sigma L9393), rat IgG2a-anti-CD31 (1:100, BD 550274), mouse IgG1-anti-VWF (1:100, Dako M0616), rabbit IgG-anti-βIIITubulin (1:200, Abcam ab246929, ab18207), mouse IgG2b-anti-βIIITubulin (1:2000, Biolegend 801201). Alexa Fluor secondary antibodies (Thermo Fisher Scientific) and Alexa Fluor αBTX (Thermo Fisher Scientific B13422, B35451, B35450) were used at 1:500 dilution.

Three-steps staining was used to avoid interaction between rat antibodies in CD31 MCAM staining: first step, CD31 staining; second step, secondary antibody for CD31; third step, MCAM staining (the MCAM antibody used in the experiments is conjugated with Alexa Fluor 647). Incubation steps and washing steps were maintained as described, therefore, some more days were needed to finish this specific staining.

Mounting procedure was specifically designed for samples with this thickness. Slides were previously prepared, strips of 4-5 layers of electrical tape were located in the edges of the slide, forming a window in the slide to accommodate muscles. At least, 1mL of mounting media was used (Fluoromount G [EMS 17984-25]), and then a coverslip was located over, paying special attention to avoid the formation of bubbles. These “mounting structures” were let completely dry at RT in a dark place in horizontal position, during approximately 7-10 days (refilling with mounting media in edges when necessary). Drying process can be shortened when using boxes hermetically closed with the “mounting structures” located over drying materials (silica gel or rice). As NMJs can be observed all around the piece of the muscle, these “mounting structures” can be improved by using two coverslips instead of using a slide and a coverslip, to allow the acquisition of images all around the surface of the sample. Once completely dry, they were stored at 4°C until confocal analysis.

### Confocal image acquisition and analysis

Images were acquired using a Zeiss LSM900 microscope (20x and 40x objectives) and analysed with ZEN 2.3 blue edition software and Arivis Vision 4D software. Each acquired image was composed of 50-100 optical sections, capturing a comprehensive Z-stack of the NMJ structure. Subsequently, each NMJ within the acquired images was individually analysed for further investigation and quantitative assessments.

All statistical analyses were performed using GraphPad Prism 6.0 software. Data are given as mean ± s.d. When counting NMJs per animal, at least 50 NMJs/animal were counted. When counting contacts of kranocytes with vessels, each prolongation was checked and followed in the 3D structure (slide by slide); at least 30 NMJs were counted per animal.

*Bona fide* tSCs have been considered as tSCs with continuous staining of S100β and MCAM that completely cover the endplate; tSCs with partial coverage of the endplate, low/absent expression of S100β and/or MCAM abnormal structure have been considered atypical tSCs

### Transmission electron microscopy and pre-embedding immunogold

Electron microscopy: For conventional electron microscopy studies, EDL muscles were fixed by immersion with 2% paraformaldehyde and 2.5% glutaraldehyde for 24 hours. Then 200 µm sections were cut on a vibratome (Leica VT-1000). Sections were post-fixed with 2% osmium tetroxide for 1.5 hours, rinsed, dehydrated and embedded in araldite (Durcupan, Fluka). Semithin sections (1.5 mm) were cut with an Ultracut UC-6 (Leica), mounted on gelatine-coated slides and stained with 1% toluidine blue. These sections were examined under a light microscope (Eclipse E200, Nikon). To identify individual cell types, ultrathin sections (60-70nm) were cut, stained with lead citrate (Reynolds solution) and examined under a transmission electron microscope (Tecnai Spirit G2, FEI). Images were acquired using Radius software (Version 2.1) with a Xarosa digital camera (EMSIS GmbH)

Immunocytochemistry: For pre-embedding staining, EDL muscles were fixed by immersion with 4% paraformaldehyde. Then 100 µm sections were cut on a vibratome (Leica VT-1000). Pre-embedding immunogold staining was performed by incubating sections in 1:25 primary antibody (rat IgG2a-anti-MCAM [BD 562230] or rat IgG2a-anti-CD34 [BD 553731]) and in the colloidal gold-conjugated secondary antibody goat anti-rat (1:50; UltraSmall, Aurion) as previously described ^84^.

Then the sections were post-fixed with 1% osmium tetroxide with 7% glucose for 30 min, rinsed, dehydrated and embedded in Araldite (Durcupan, Fluka). Semithin sections (1.5 mm) were cut with an Ultracut UC-6 (Leica), mounted on gelatine-coated slides. These sections were examined under a light microscope (Eclipse E200, Nikon), previously to stained with 1% toluidine blue, to identify positive cells. Then consecutive semithin sections with positive cells were re-embedded, and ultrathin sections (60-70 nm) were cut. These ultrathin sections were stained with lead citrate (Reynolds solution) and examined under a transmission electron microscope (Tecnai Spirit G2, FEI). Images were acquired using Radius software (Version 2.1) with a Xarosa digital camera (EMSIS GmbH).

### Dataset processing and scRNAseq analysis

Published data bases were used ^20,32,47,48^. Each dataset has been processed using a common pipeline, run with Scanpy. Cells with fewer than a set number of expressed genes was removed, ranging between 1 and 250, depending on the dataset. Datasets were normalised using the scanpy.pp.normalize_per_cell function, and log1p-transformed. Feature selection was performed using triku ^85^. Dimensionality reduction was performed using Principal Component Analysis (PCA) with 30 components. Then, the k-Nearest Neighbor (kNN) graph was computed using the PCA matrix, with cosine similarity. In datasets with different samples, the kNN graph was constructed using bbknn^86^. For dataset visualisation, UMAP (Uniform Manifold Approximation and Projection) ^87^ was used. Clustering was performed using leiden ^88^. Gene Ontology (GO) analysis was performed in data set D ^20^.

### Cell type and kranocyte assignment in scRNAseq analysis

In order to assign cell types and kranocytes to the different datasets, a dictionary with markers and cell types was mapped using a custom cell assignment algorithm (https://github.com/alexmascension/cell_asign).

For major cell types in Figure 1q and Supplementary Figure 1m and n were FAP Lum+ (*Apod, Lum, Ly6a, Pdgfra, Mfap5, Dcn*), FAP Prg4+ (*Prg4, Fbn1, Ly6a, Pdgfra, Mfap5, Dcn*), Endothelial (*Pecam1, Kdr, Fabp4, Cav1, Cdh5, Tek*), Tenocyte (*Scx, Tnmd, Mkx, Col12a1, Col1a1, Tnc, Fmod, Comp*), Glial cell (*Plp1, Kcna1, S100b, Mbp, Mpz*), Pericyte (*Rgs5, Notch3, Myl9, Ndufa4l2, Itga7, Myh11, Pln, Abcc9*), Myonuclei (*Tnnc2, Myh4, Acta1, Ckm, Tpm2, Eno3, Slc25a4*), Satellite cell (*Pax7, Myod1, Chodl, Vcam1, Sdc4, Myf5*), Immune (*H2-Aa, Cd74*), Monocyte (*Csf1r, Adgre1*), APC (*H2-Eb1, H2-Ab1*), B cell (*Cd19, Cd22, Ms4a1, Ptprc*), T cell (*Cd3d, Cd3e, Cd3g, Cd8a, Cd4, Ptprc, Cd28*), Macrophage (*Itgam, Csf1r, Adgre1, Itgb1, Cd68*), Neutrophil (*S100a8, S100a9, Itgam, Cd14*).

For kranocytes in Figure 1q and Supplementary Figure 1m, the markers used for the assignment were *6030408B16Rik, Smim41, Col9a2, Dlk1, Shisa3, Saa1, Nipal1*.

For Schwann cells in Figure 1b, the markers used were: for SCs (*S100b, Plp1, Sox10, Kcna1*), for mSCs (*Mpz, Prx, Gatm, Mal, Cldn19, Mal, Pllp, Mt3, Bcas1, Emid1, Mag, Grp, Ptpre, Kcnk1, Plekha4, Ugt8a*), and for tSCs (*Mcam, Ng2, Ptn, Cadm1, Col20a1, Lgi4, Postn, Nrn1, Ncam1, Tinagl1, Sfrp1, Matn4, Acsbg1, Lyz2, Gm2115, Chl1, Bche, Nrxn1, Ptprz1*). In this cluster of tSCs, we cannot elude the inclusion of nmSCs, what will need specific future analysis.

Notebooks used for dataset processing and analysis are available at https://github.com/alexmascension/kranocyte-analysis.

