## Supplementary info for "Unveiling the neuro-vascular interplay in the skeletal muscle in health, injury and disease"

### SUPPLEMENTARY FIGURES

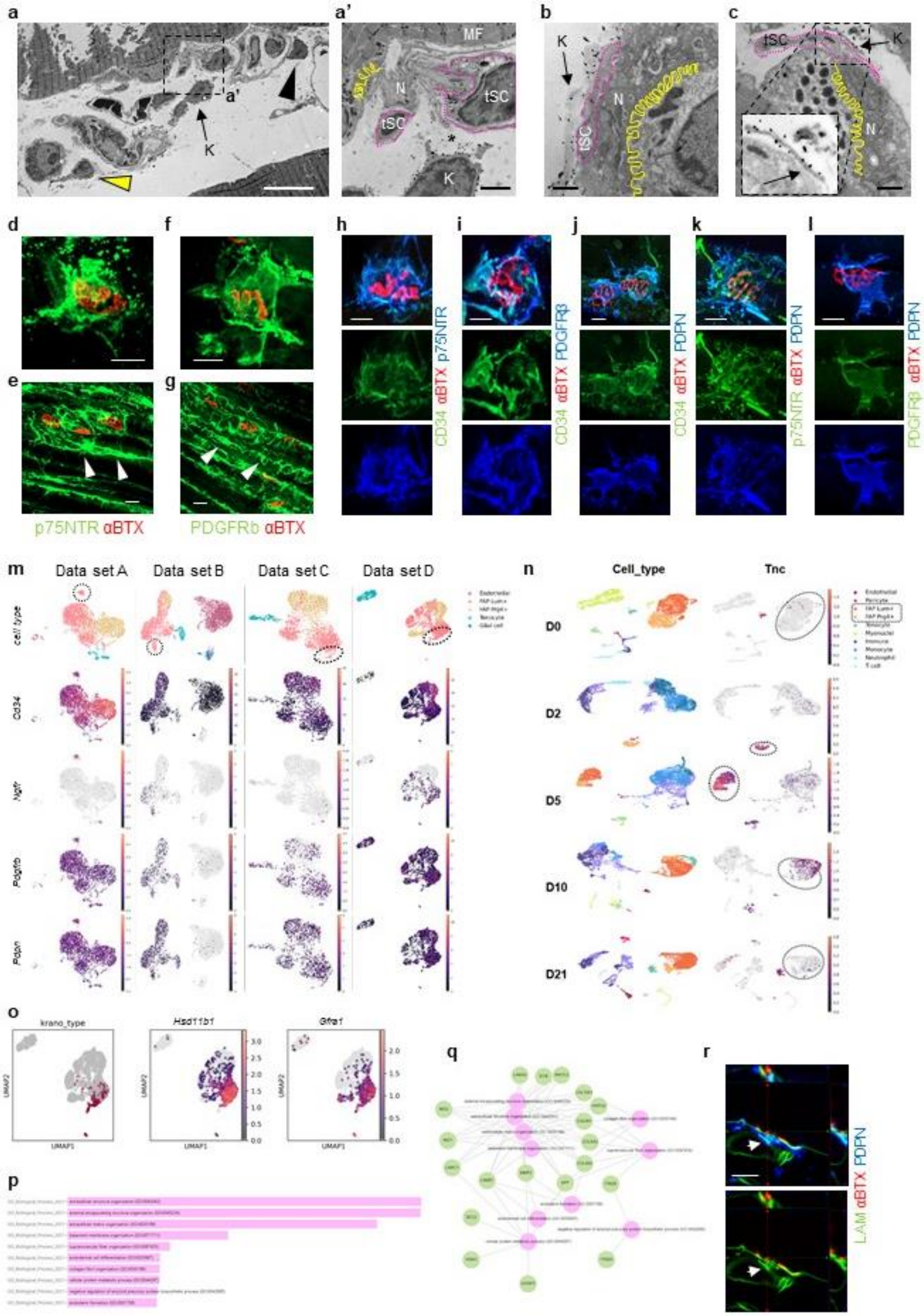

Supplementary Figure 1. **Cell markers, scRNAseq and GO analyses for kranocytes. a** TEM analysis of muscle interstitial cells, CD34+ cells are shown (kranocyte, arrow; CD34+ perivascular cell, black arrowhead; CD34+ interstitial cell, yellow arrowhead); **a’** is an insert of **a**, area between the tSC and the kranocyte is shown (asterisk). **b, c** TEM analysis of terminal Schwann cell (tSC) and kranocyte localization. **d, e** Confocal images of p75NTR, showing a kranocyte (**d**) and also other interstitial cells (**e**, arrowheads). **f, g** Confocal images of PDGFRβ, showing a kranocyte (**f**) and also other interstitial cells (**g**, arrowheads). **h** Confocal image of CD34 and p75NTR. **i** Confocal images of CD34 and PDGFRβ. **j** Confocal images of CD34 and Podoplanin (PDPN). **k** Confocal images of PDPN and p75NTR. **l** Confocal images of PDPN and PDGFRβ. **m** UMAP plots of scRNAseq analysis of data sets A [1], B [2], C [3], D [4]: *Cd34*, *Ngfr* and *Pdgfrb* and *Pdpn* gene expression are shown, kranocytes clusters are highlighted (dotted lines). **n** UMAP plots of scRNAseq analysis of data set C [3] after injury, expression of the cell adhesion molecule tenascin C (*Tnc*) is shown; FAP clusters are highlighted (dotted circles). **o** UMAP plots of scRNAseq analysis of data set D [4], colocalization of kranocytes cluster (first panel) and Hsd11b1+ Gfra1+ population (second and third panels) (expression of *Hsd11b1* and *Gfra1* are shown). **p, q** Gene Ontology (GO) analysis of data set D [4]; bar graph showing the most relevant biological processes (**p**) and network graph showing the genes involved in the biological processes in **p** (**q**). **r** Confocal image and orthogonal projections of Podoplanin (PDPN) and Laminin (LAM) over an endplate, colocalization of PDPN and LAM is shown (arrows). Scale bar: 50 µm (**e**, **g**), 20 µm (**d**, **f**, **h**, **i**, **j**, **k**, **l**, **r**), 10 µm (**a**), 2 µm (**a’**), 1 µm (**b**, **b’**). Endplate was labelled with αBungarotoxin (αBTX) for acetylcholine receptors (AChRs). Endplates are highlighted by white solid lines. TEM analysis: immunogold staining of CD34. TEM image: endplate (yellow line), tSC (pink dotted line), capillary (C), kranocyte (K), muscle fibre (MF), terminal nerve (N), terminal Schwann cell (tSC).

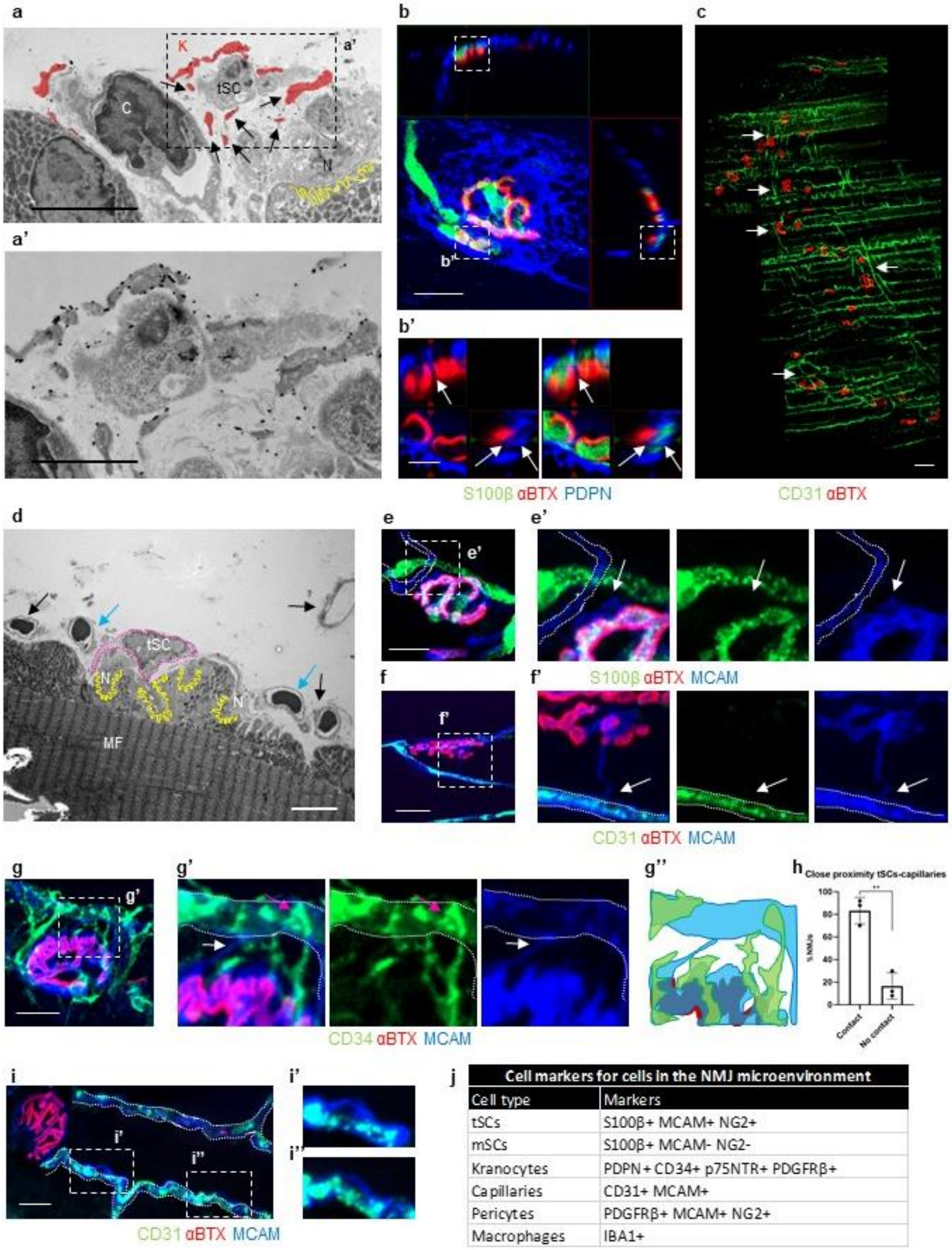


| Cell markers for cells in the NMJ microenvironment | |
|---|---|
| Cell type | Markers |
| tSCs | S100β+ MCAM+ NG2+ |
| mSCs | S100β+ MCAM- NG2- |
| Kranocytes | PDPN+ CD34+ p75NTR+ PDGFRβ+ |
| Capillaries | CD31+ MCAM+ |
| Pericytes | PDGFRβ+ MCAM+ NG2+ |
| Macrophages | IBA1+ |

Supplementary Figure 2. **tSCs and kranocytes interactions with microenvironment. a** TEM analysis of a NMJ, where a kranocyte (colored in red) and its protrusions under the terminal Schwann cell (tSC)-body (kranocyte colored in red, arrows) are shown; **a’** is an insert of **a**. **b** Confocal image and orthogonal projections of S100β and Podoplanin (PDPN); **b’** are inserts of **b** where small protrusions of PDPN are located under S100β staining (arrows). **c** Tile confocal images of CD31 in the vicinity of NMJs, branching points are shown (white arrows). **d** TEM analysis of vascularization in the vicinity of a NMJ: several capillaries are observed (arrows), two of them in close proximity to NMJ-capping cells and synaptic area (blue arrows). **e, e'** Confocal image of S100β and MCAM, where MCAM+S100β- protrusion (white arrow) from a tSC is shown; **e'** is an insert of **e**. **f, f'** Confocal image of MCAM and CD31, where a sprout MCAM+ from a tSC contact a capillary (white arrow); **f'** is an insert of **f**. **g** Confocal image of MCAM and CD34 showing tSCs and kranocytes; **g'** is an insert of **g**, showing a sprout from the tSC (white arrow) and a sprout from the kranocytes (end feet structure, pink arrow) both of them reaching the capillary; **g''** schematic representation of **g'**. **h** Quantification of NMJs in close proximity to capillaries through their tSCs. **i** Confocal images of MCAM and CD31, where MCAM+CD31+ capillaries (**i**, white dotted lines) and pericites (**i', i''**) are shown; **i'** and **i''** are inserts of **i**. **j** Summary table of the markers used to identify each cell type. Scale bar: 50 µm (**c**), 20 µm (**b**, **e**, **f**, **g**, **i**), 5 µm (**a**, **b'**, **d**), 2 µm (**a'**). Endplate was labelled with αBungarotoxin (αBTX) for acetylcholine receptors (AChRs). Capillaries are highlighted by white dotted lines (**e**, **e'**, **f'**, **g'**, **i**). TEM analysis: immunogold staining of CD34. TEM image: endplate (yellow line), tSC (pink dotted line), capillary (C), kranocyte (K), muscle fibre (MF), terminal nerve (N), terminal Schwann cell (tSC). Statistics Student's unpaired T-test was used for comparisons: ns, not significant, $p > 0.05$, *$p<0.05$, **$p \leq 0.01$, ***$p \leq 0.001$, ****$p \leq 0.0001$. Quantifications show mean ± s.d.

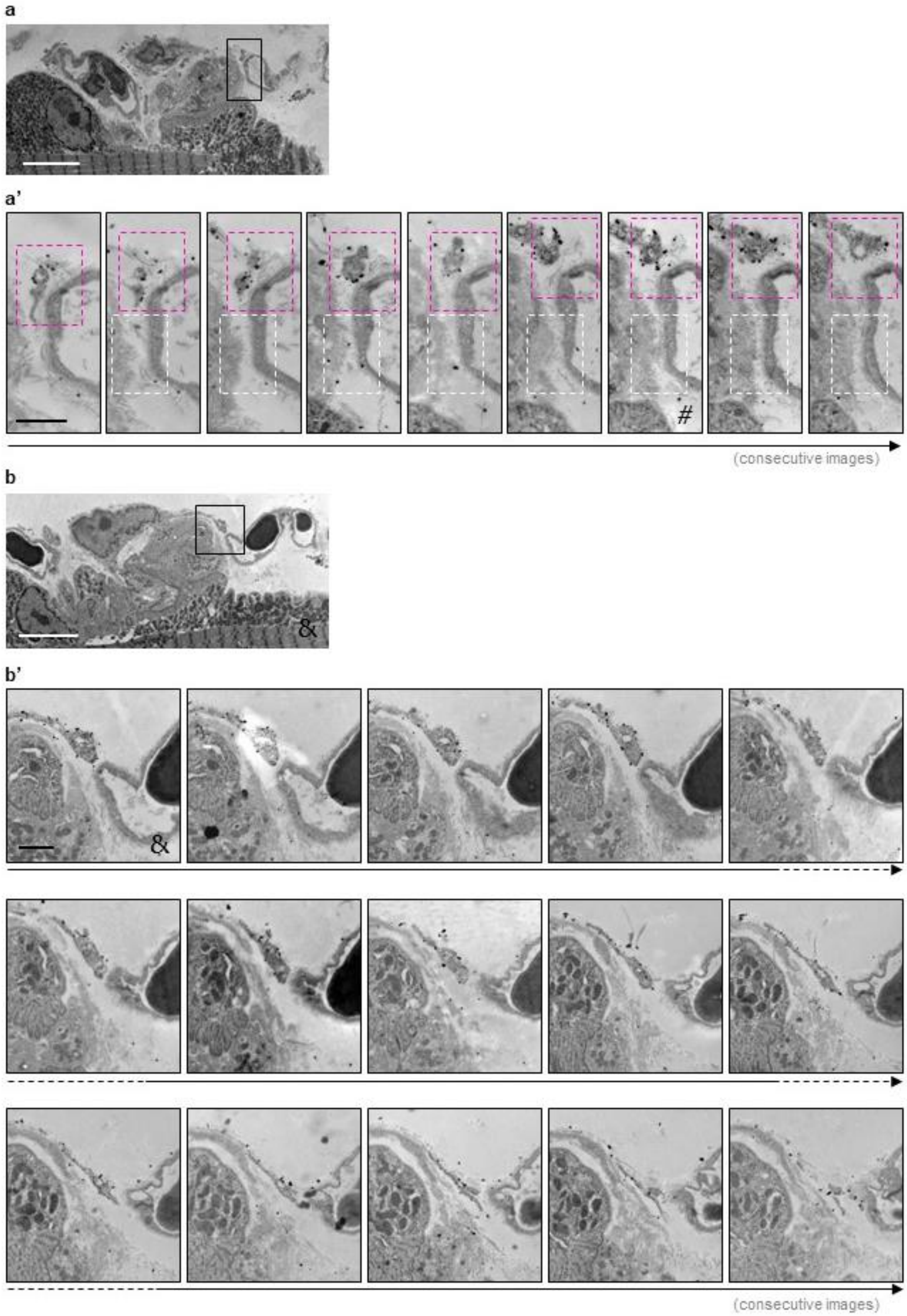


Supplementary Figure 3. **Sequential images of NMJ-capping cells reaching a capillary.** Sequential images of TEM analysis of a NMJ where it is observed a tSC protrusion towards a capillary (**a, a’**, white dotted square) and sprouts of a kranocyte spreading towards a capillary (**a, a’**, pink dotted square and **b, b’**). Scale bar: 5 µm (**a**, **b**), 1 µm (**a’**, **b’**). TEM analysis: immunogold staining of CD34. Characters indicate coincident images with figure 2 (#, &).

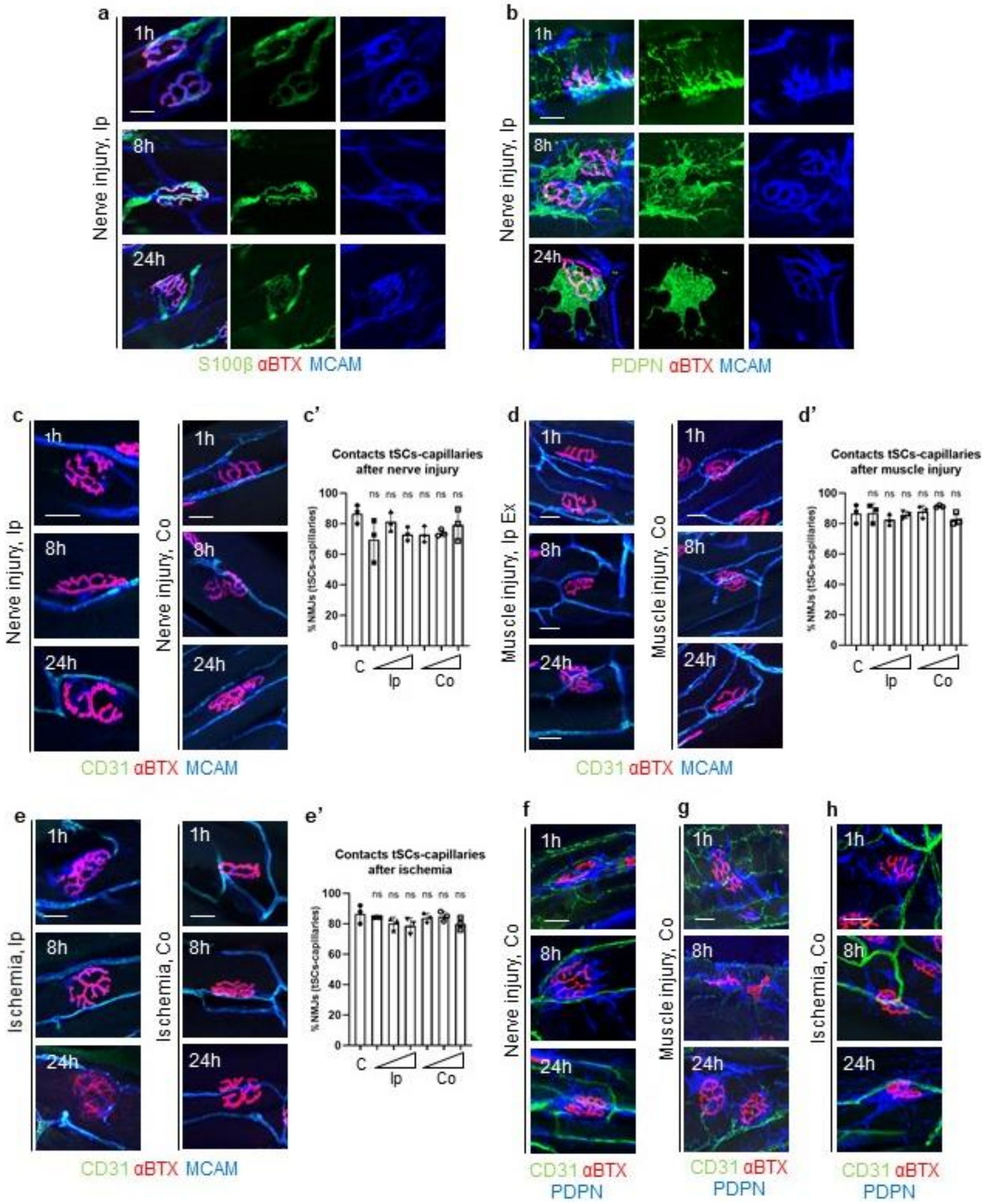


Supplementary Figure 4. **tSC-capillaries interaction and contralateral muscles. a, b** Confocal images of MCAM and S100β (**a**) or Podoplanin (PDPN) (**b**) after nerve injury in ipsilateral (Ip) muscles, showing no detectable structural changes in terminal Schwann cells (tSCs). **c, d. e** Confocal images of MCAM and CD31 after nerve injury (**c**), muscle injury (**d**) and ischemia (**e**) in Ip and contralateral (Co) muscles. **c', d', e'** Quantification of contacts between tSCs and capillaries after nerve injury (**c'**), muscle injury (**d'**) and ischemia (**e'**) in Ip and Co muscles. **f-h** Confocal images of PDPN and CD31 after nerve injury (**f**), muscle injury (**g**) and ischemia (**h**) in Co muscles. Scale bar: 20 µm (**a**, **b**, **c**, **d**, **e**, **f**, **g**, **h**, **i**). Endplate was labelled with αBungarotoxin (αBTX) for acetylcholine receptors (AChRs). Statistics Student's unpaired T-test was used for comparisons: ns, not significant, $p > 0.05$, $^{*}p<0.05$, $^{**}p \leq 0.01$, $^{***}p \leq 0.001$, $^{****}p \leq 0.0001$. Quantifications show mean ± s.d.

### Supplementary tables

Supplementary Table 1. **DEGs of mSCs and tSCs**. DEGs obtained from scRNAseq analysis of muscle SCs [1]: mSCs (first column), tSCs (second column). *p*-values are shown. Expression of *Mcam* in tSCs is highlighted in yellow.

Supplementary Table 2. **DEGs of kranocytes**. DEGs obtained from scRNAseq analysis of muscle interstitial cells: data set A [1], B [2], C [3] and D [4]. For each dataset, DEGs of kranocytes against the rest of cells were calculated using scanpy's *scanpy.tl.rank_genes_groups* function. Each table shows the ranking of the top 100 DEGs, as well as their p-values and expression log fold changes. Expression of *Hsd11b1* (green), *Lama2, Lamb1* and *Lamc1* (blue) and *Gfra1* and *Gfra2* (orange) are highlighted.

Supplementary Table 3. **Enrichment table of Gene Ontology analysis of kranocytes**. GO Biological Process analysis of data set D [4]. p-values, q-values, z-scores and combined scores are shown.

### Supplementary videos

Supplementary Video 1. **Close proximity between tSCs and capillaries**. tSCs (S100β, blue) and capillaries (CD31, green) are shown. Endplate (red) was labelled with αBungarotoxin (αBTX) for acetylcholine receptors (AChRs).

Supplementary Video 2. **tSC-sprout towards a capillary_MCAM+CD31-**. A tSC (MCAM, blue) and capillaries (CD31, green) are shown. A tSC-sprout (MCAM+ CD31-) is observed. Endplate (red) was labelled with αBungarotoxin (αBTX) for acetylcholine receptors (AChRs).

Supplementary Video 3. **Kranocytes contacting capillaries**. Kranocytes (Podoplanin, PDPN, blue) and capillaries (CD31, green) are shown. Endplate (red) was labelled with αBungarotoxin (αBTX) for acetylcholine receptors (AChRs).
